# Zygotic contractility awakening during mouse preimplantation development

**DOI:** 10.1101/2021.07.09.451745

**Authors:** Özge Özgüç, Ludmilla de Plater, Varun Kapoor, Anna Francesca Tortorelli, Jean-Léon Maître

## Abstract

Actomyosin contractility is a major engine of preimplantation morphogenesis, which starts at the 8-cell stage during mouse embryonic development. Contractility becomes first visible with the appearance of periodic cortical waves of contraction (PeCoWaCo), which travel around blastomeres in an oscillatory fashion. How contractility of the mouse embryo becomes active remains unknown. We have taken advantage of PeCoWaCo to study the awakening of contractility during preimplantation development. We find that PeCoWaCo become detectable in most embryos only after the 2^nd^cleavage and gradually increase their oscillation frequency with each successive cleavage. To test the influence of cell size reduction during cleavage divisions, we use cell fusion and fragmentation to manipulate cell size across a 20-60 *μ*m range. We find that the stepwise reduction in cell size caused by cleavage divisions does not explain the presence of PeCoWaCo or their accelerating rhythm. Instead, we discover that blastomeres gradually decrease their surface tensions until the 8-cell stage and that artificially softening cells enhances PeCoWaCo prematurely. Therefore, during cleavage stages, cortical softening awakens zygotic contractility before preimplantation morphogenesis.

## Introduction

During embryonic development, the shape of animal cells and tissues largely relies on the contractility of the actomyosin cortex (Coravos et al., 2017; Murrell et al., 2015; Özgüç and Maître, 2020). The actomyosin cortex is a sub-micron-thin layer of crosslinked actin filaments, which are put under tension by non-muscle myosin II motors (Kelkar et al., 2020). Tethered to the plasma membrane, the actomyosin cortex is a prime determinant of the stresses at the surface of animal cells (Clark et al., 2014; Kelkar et al., 2020). Contractile stresses of the actomyosin cortex mediate crucial cellular processes such as the ingression of the cleavage furrow during cytokinesis (Fujiwara and Pollard, 1976; Straight et al., 2003; Yamamoto et al., 2021), the advance of cells’ back during migration (Eddy et al., 2000; Tsai et al., 2019) or the retraction of blebs (Charras et al., 2006; Taneja and Burnette, 2019). At the tissue scale, spatiotemporal changes in actomyosin contractility drive apical constriction (Martin et al., 2009; Solon et al., 2009) or the remodeling of cell-cell contacts (Bertet et al., 2004; Maitre et al., 2012). Although tissue remodeling takes place on timescales from tens of minutes to hours or days, the action of the actomyosin cortex is manifest on shorter timescales of tens of seconds (Coravos et al., 2017; Michaud et al., 2021; Özgüç and Maître, 2020). In fact, actomyosin is often found to act via pulses of contraction during morphogenetic processes among different animal species from nematodes to human cells (Baird et al., 2017; Bement, 2015; Blanchard et al., 2010; Kim and Davidson, 2011; Maître et al., 2015; Martin et al., 2009; Munro et al., 2004; Roh-Johnson et al., 2012; Solon et al., 2009). A pulse of actomyosin begins with the polymerization of actin filaments and the sliding of myosin mini-filaments until maximal contraction of the local network within about 30 s (Dehapiot et al., 2020; Ebrahim et al., 2013; Martin et al., 2009). Then, the actin cytoskeleton disassembles, and myosin is inactivated, which relaxes the local network for another 30 s (Mason et al., 2016; Munjal et al., 2015; Vasquez et al., 2014). These cycles of contractions and relaxations are governed by the turnover of the Rho GTPase and its effectors, which are well-characterized regulators of actomyosin contractility (Bement, 2015; Graessl et al., 2017; Munjal et al., 2015). In instances where a sufficient number of pulses occur, pulses of contraction display a clear periodicity. The oscillation period of pulsed contractions ranges from 60 to 200 s (Baird et al., 2017; Bement, 2015; Maître et al., 2015; Solon et al., 2009). The period appears fairly defined for cells of a given tissue but can vary between tissues of the same species. What determines the oscillation period of contraction is poorly understood, although the Rho pathway may be expected to influence it (Bement, 2015; Munjal et al., 2015; Vasquez et al., 2014). Finally, periodic contractions can propagate into travelling waves. Such periodic cortical waves of contraction (PeCoWaCo) were observed in cell culture, starfish, and frog oocytes as well as in mouse preimplantation embryos (Bement, 2015; Driscoll et al., 2015; Kapustina et al., 2013; Maître et al., 2015). In starfish and frog oocytes, mesmerizing Turing patterns of Rho activation with a period of 80 s and a wavelength of 20 *μ*m appear in a cell cycle dependent manner (Bement, 2015; Wigbers et al., 2021). Interestingly, experimental deformation of starfish oocytes revealed that Rho activation wave front may be coupled to the local curvature of the cell surface (Bischof et al., 2017), which was proposed to serve as a mechanism for cells to sense their shape (Wigbers et al., 2021). In mouse embryos, PeCoWaCo with a period of 80 s were observed at the onset of blastocyst morphogenesis (Maître et al., 2015; Maître et al., 2016). What controls the propagation velocity, amplitude, and period of these waves is unclear and the potential role of such evolutionarily conserved phenomenon remains a mystery.

During mouse preimplantation development, PeCoWaCo become visible before compaction (Maître et al., 2015), the first morphogenetic movements leading to the formation of the blastocyst (Maître, 2017; Özgüç and Maître, 2020; White et al., 2018). During the second morphogenetic movement, prominent PeCoWaCo are displayed in prospective inner cells before their internalization (Maître et al., 2016). In contrast, cells remaining at the surface of the embryo display PeCoWaCo of lower amplitude due to the presence of a domain of apical material that inhibits the activity of myosin (Maître et al., 2016). Then, during the formation of the blastocoel, high temporal resolution time-lapse hint at the presence of PeCoWaCo as microlumens coarsen into a single lumen (Dumortier et al., 2019). Therefore, PeCoWaCo appear throughout the entire process of blastocyst formation (Özgüç and Maître, 2020). However, little is known about what initiates and regulates PeCoWaCo. The analysis of maternal zygotic mutants suggests that PeCoWaCo in mouse blastomeres result primarily from the action of the non-muscle myosin heavy chain IIA (encoded by *Myh9*) rather than IIB (encoded by *Myh10*) (Schliffka et al., 2021). Dissociation of mouse blastomeres indicates that PeCoWaCo are cell-autonomous since they persist in single cells (Maître et al., 2015). Interestingly, although removing cell-cell contacts free-up a large surface for the contractile waves to propagate, the oscillation period seems robust to the manipulation (Maître et al., 2015). Similarly, when cells form an apical domain taking up a large portion of the cell surface, the oscillation period does not seem to be different from cells in which the wave can propagate on the entire cell surface (Maître et al., 2016). This raises the question of how robust PeCoWaCo are to geometrical parameters, especially in light of recent observations in starfish oocytes (Bischof et al., 2017; Wigbers et al., 2021). This question becomes particularly relevant when considering that, during preimplantation development, cleavage divisions halve cell volume with each round of cytokinesis (Aiken et al., 2004; Niwayama et al., 2019).

In this study, we investigate how the contractility of the cleavage stages emerges before initiating blastocyst morphogenesis. We take advantage of the slow development of the mouse embryo to study thousands of pulsed contractions and of the robustness of the mouse embryo to size manipulation to explore the geometrical regulation of PeCoWaCo. We discover that the initiation, maintenance, or oscillatory properties of PeCoWaCo do not depend on cell size. Instead, we discover a gradual softening of blastomeres with each successive cleavage, which conceals PeCoWaCo. Together, this study reveals how preimplantation contractility is robust to the geometrical changes of the cleavage stages during which the zygotic contractility awakens.

## Results

### PeCoWaCo during cleavage stages

PeCoWaCo have been observed at the 8-, 16-cell, and blastocyst stages. To know when PeCoWaCo first appear, we imaged embryos during the cleavage stages and performed Particle Image Velocimetry (PIV) and Fourier analyses (Fig 1A-C, Movie 1). This reveals that PeCoWaCo are detectable in fewer than half of zygote and 2-cell stage embryos and become visible in most embryos from the 4-cell stage onwards (Fig 1D). Furthermore, PeCoWaCo only display large amplitude from the 4-cell stage onwards (Fig 1B-C). Interestingly, the period of oscillations of the detected PeCoWaCo shows a gradual decrease from 200 s to 80 s between the zygote and 8-cell stages (Fig 1E). The acceleration of PeCoWaCo rhythm could simply result from the stepwise changes of cell size after cleavage divisions. Indeed, we reasoned that if the contractile waves travel at constant velocity, the period will scale with cell size and shape. This is further supported by the fact that PeCoWaCo are detected at the same rate and with the same oscillation period during the early or late halves of the 2-, 4- and 8-cell stages (Fig S1). Therefore, we set to investigate the relationship between cell size and periodic contractions.

**Figure 1:**
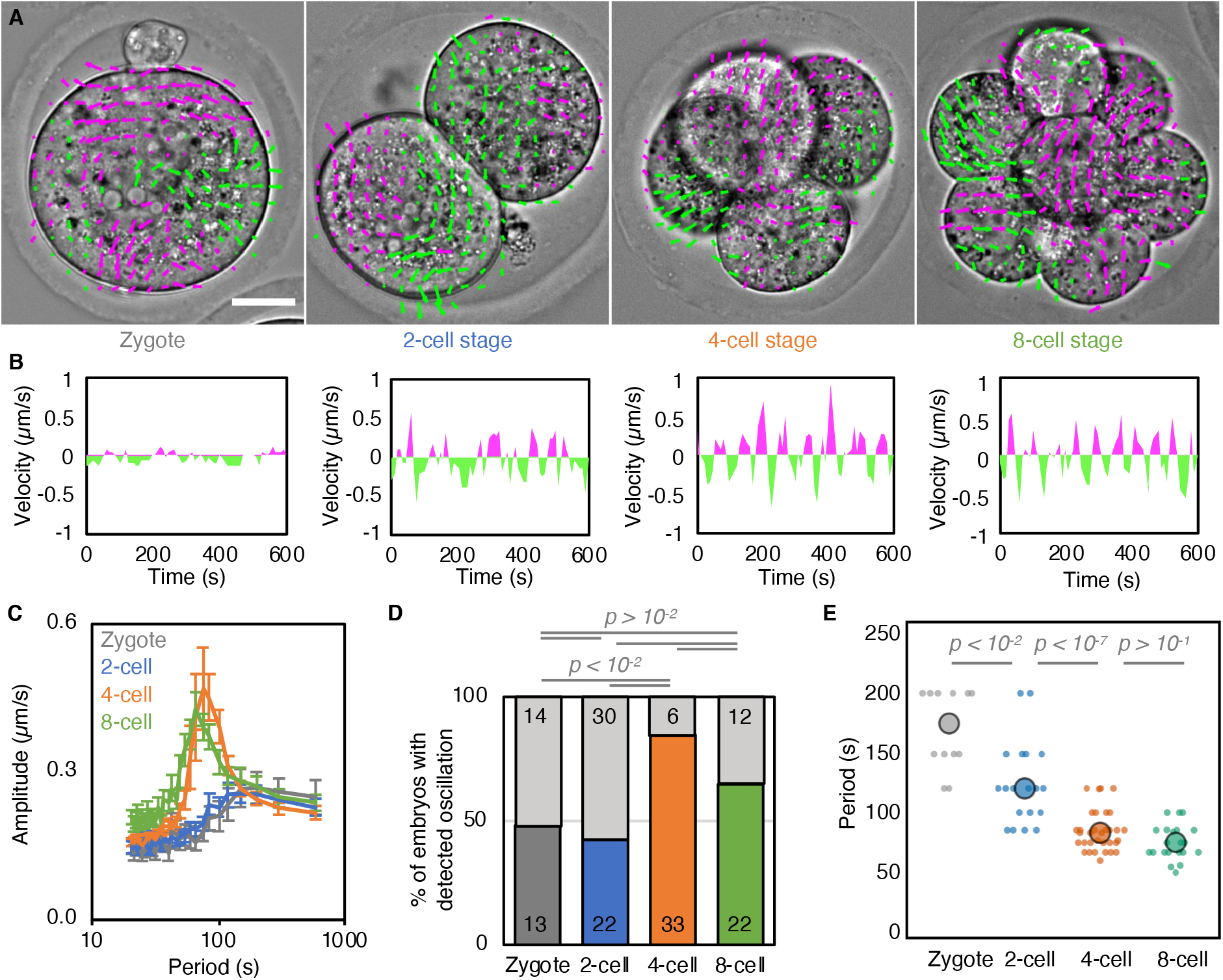
Analysis of PeCoWaCo during cleavage stages. **A)** Representative images of a short-term time-lapse overlaid with a subset of velocity vectors from Particle Image Velocimetry (PIV) analysis during cleavage stages (Movie 1). Magenta for positive and green for negative Y directed movement. Scale bars, 20 *μ*m. **B)** Velocity over time for a representative velocity vector of each embryo shown in A. **C)** Mean power spectrum resulting from Fourier transform of PIV analysis of Zygote (grey, n = 13), 2-cell (blue, n = 22), 4-cell (orange, n = 33) and 8-cell (green, n = 22) stages embryos showing detectable oscillations. Data show as mean ± SEM. **D)** Proportion of Zygote (grey, n = 27), 2-cell (blue, n = 52), 4-cell (orange, n = 39), 8-cell stage (green, n = 34) embryos showing detectable oscillations after Fourier transform of PIV analysis. Light grey shows non-oscillating embryos. Chi^2^ p values comparing different stages are indicated. **E)** Oscillation period of Zygote (grey, n = 13), 2-cell (blue, n = 22), 4-cell (orange, n = 33), 8-cell (green, n = 22) stages embryos. Larger circles show median values. Student’s t test p values are indicated.

### Cell size is not critical for the initiation or maintenance of PeCoWaCo

First, to test whether the initiation of PeCoWaCo in most 4-cell stage embryos depends on the transition from the 2- to 4-cell stage blastomere size, we prevented cytokinesis. Using transient exposure to Vx-680 to inhibit the activity of Aurora kinases triggering chromosome separation, we specifically blocked the 2- to 4-cell stage cytokinesis without compromising the next cleavage to the 8-cell stage (Fig 2A-B, Movie 2). This causes embryos to reach the 4-cell stage with tetraploid blastomeres the size of 2-cell stage blastomeres. At the 4-cell stage, we detect PeCoWaCo in most embryos whether they have 4- or 2-cell stage size blastomeres (Fig 2C). Furthermore, the period of oscillation is identical to 4-cell stage embryos in both control and drug-treated conditions (Fig 2D). This suggests that 4-cell stage blastomere size is not required to initiate PeCoWaCo in the majority of embryos.

**Figure 2:**
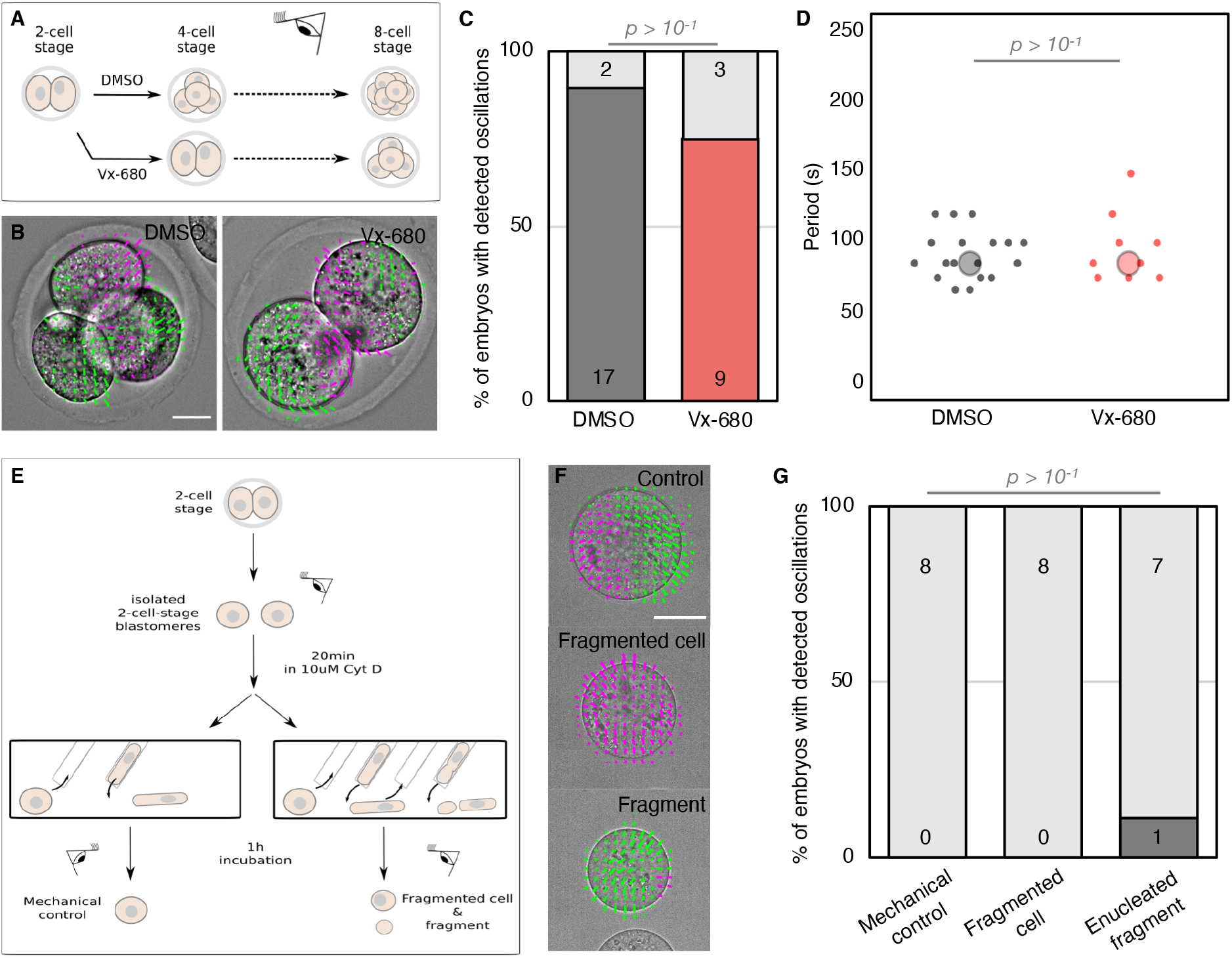
Initiation of PeCoWaCo is independent of cell size. **A)** Schematic diagram of PeCoWaCo analysis after blocking the 2^nd^ cleavage division with 2.5 *μ*m Vx680. **B)** Representative images of DMSO and Vx-680 treated embryos overlaid with a subset of velocity vectors from Particle Image Velocimetry (PIV) analysis (Movie 2). **C-D)** Proportion (C) of embryos showing detectable oscillations and their detected period (D, DMSO n = 19 and Vx-680 n = 12). Chi^2^ (C) and Student’s t test (D) p values comparing two conditions are indicated. Larger circles show median values. **E)** Schematic diagram of PeCoWaCo analysis after fragmentation of 2-cell stage blastomeres. **F)** Representative images of mechanical control, fragmented cell and enucleated fragments overlaid with a subset of velocity vectors from PIV analysis (Movie 3). **G)** Proportion of cells showing detectable oscillations in mechanical controls (n = 8), fragmented cells (n= 8) and enucleated fragments (n=9). Fisher exact test p values comparing different conditions are indicated. Light grey shows non-oscillating embryos.

Then, we tested whether PeCoWaCo could be triggered prematurely by artificially reducing 2-cell stage blastomeres to the size of a 4-cell stage blastomere. First, we analyzed for the presence of PeCoWaCo in dissociated 2-cell stage blastomeres and then proceeded to reduce their size (Fig 2E-F, Movie 3). To reduce cell size, we treated dissociated 2-cell stage blastomeres with the actin cytoskeleton inhibitor Cytochalasin D before deforming repeatedly them into a narrow pipette (Fig 2E, Fig S2). By adapting the number of aspirations of softened blastomeres, we could carefully fragment blastomeres while keeping their sister cell mechanically stressed but intact. While the fragmented cell was reduced to the size of a 4-cell stage blastomere, both fragmented and manipulated cells eventually succeeded in dividing to the 4-cell stage. After waiting 1 h for cells to recover from this procedure, we examined for the presence of PeCoWaCo. At the 2-cell stage, PeCoWaCo were rarely detected in either control or fragmented cells (Fig 2G). This suggests that 4-cell stage blastomere size is not sufficient to trigger PeCoWaCo in the majority of embryos.

### Cell size does not influence the properties of PeCoWaCo

The transition from 2- to 4-cell stage blastomere size is neither required nor sufficient to initiate PeCoWaCo. Nevertheless, the decrease in period of PeCoWaCo remarkably scales with the stepwise decrease in blastomere size (Fig 1E). Given a constant propagation velocity, PeCoWaCo may reduce their period according to the reduced distance to travel around smaller cells. To test whether cell size determines PeCoWaCo oscillation period, we set out to manipulate cell size over a broad range. By fusing varying numbers of 16-cell stage blastomeres, we built cells equivalent in size to 8-, 4- and 2-cell stage blastomeres (Fig 3A-D, Movie 4). In addition, by fragmenting 16-cell stage blastomeres, we made smaller cells equivalent to 32-cell stage blastomeres (Fig 3E-G, Movie 5)(Niwayama et al., 2019). Together, we could image 16-cell stage blastomeres with sizes ranging from 10 to 30 *μ*m in radius (Fig 3H-I). Finally, to identify how the period may scale with cell size by adjusting the velocity of the contractile wave, we segmented the outline of cells to compute the local curvature, which, unlike PIV analysis, allows us to track contractile waves and determine their velocity in addition to their period (Fig 3A) (Maître et al., 2015; Maître et al., 2016). We find that fused and fragmented 16-cell stage blastomeres show the same period, regardless of their size (Fig 3H). This could be explained if the wave velocity would scale with cell size. However, we find that the wave velocity remains constant regardless of cell size (Fig 3I). Therefore, both the oscillation period and wave velocity are properties of PeCoWaCo that are robust to changes in cell size and associated curvature.

**Figure 3:**
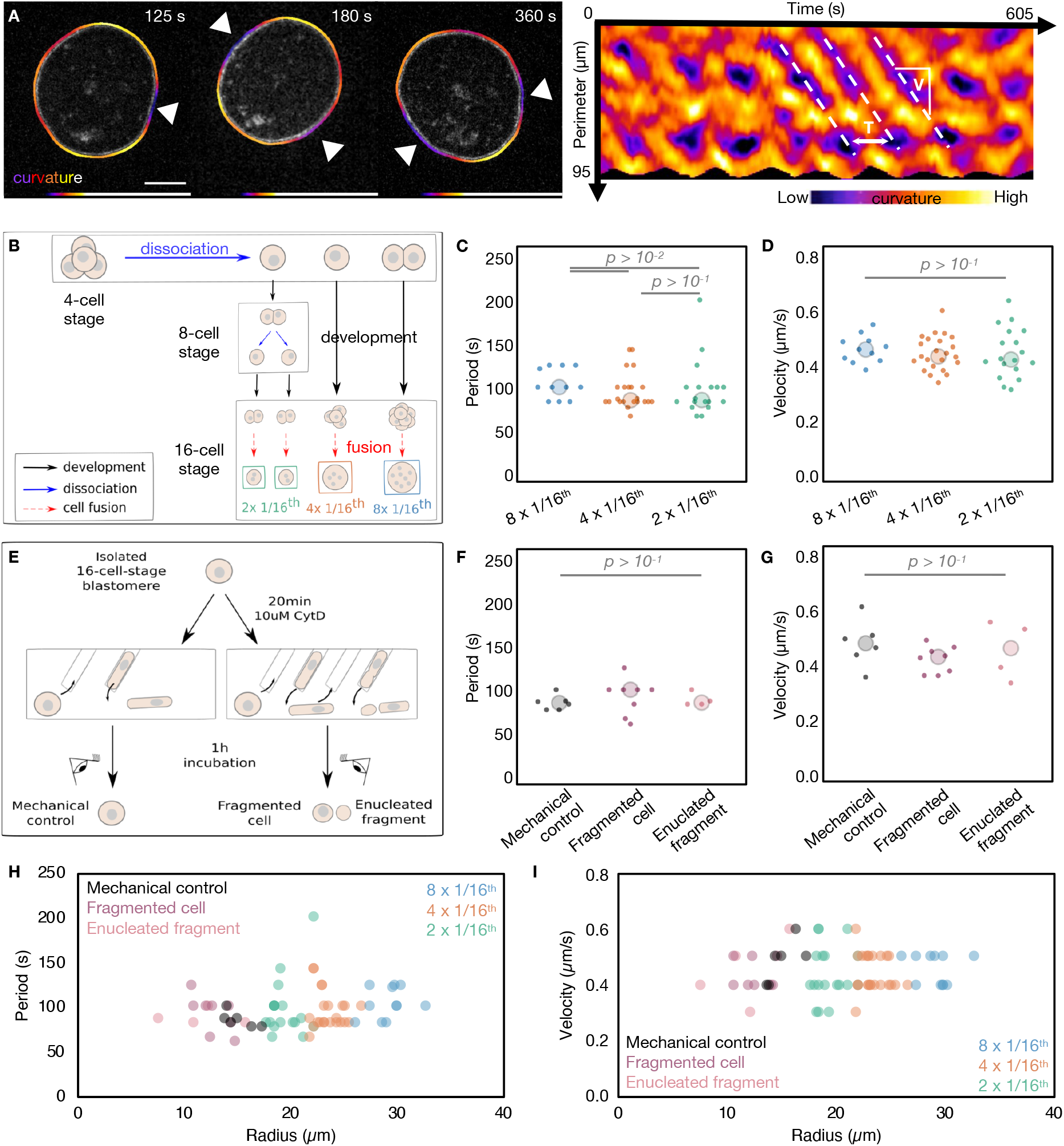
Period and velocity of PeCoWaCo are stable across a broad range of cell sizes. **A)** Surface deformation tracking for period detection and velocity measurements. Scale bar, 10 *μ*m. **B)** Schematic diagram of fusion of 16-cell stage blastomeres. **C-D)** Oscillation period (C) and wave velocity (D) of fused blastomeres. 8 × 1/16^th^ (blue, n=6), 4 × 1/16^th^ (orange, n = 12) and 2 × 1/16^th^ (green, n=7) fused blastomeres are shown. Large circles show median. **E)** Schematic diagram of fragmentation of 16-cell stage blastomeres. **F-G)** Oscillation period (F) and wave velocity (G) of fragmented blastomeres. Control (black, n = 6), fragmented cell (magenta, n= 8), enucleated fragment (pink, n=4) are shown. **H-I)** Oscillation period (H) and wave velocity (I) for size-manipulated 16-cell stage blastomeres. Larger circles show median values. Student’s t-test p values are indicated.

Fusion of cells causes blastomeres to contain multiple nuclei, while cell fragmentation creates enucleated fragments. Interestingly, enucleated fragments continued oscillating with the same period and showing identical propagation velocities as the nucleus-containing fragments (Fig 3F). These measurements indicate that PeCoWaCo are robust to the absence or presence of single or multiple nuclei and their associated functions.

Together, using fusion and fragmentation of cells, we find that PeCoWaCo oscillation properties are robust to a large range of size perturbations. Therefore, other mechanisms must be at play to regulate periodic contractions during preimplantation development.

### Cortical maturation during cleavage stages

Despite the apparent relationship between cell size and PeCoWaCo during preimplantation development, our experimental manipulations of cell size reveal that PeCoWaCo are not influenced by cell size. PeCoWaCo result from the activity of the actomyosin cortex, which could become stronger during cleavage stages and make PeCoWaCo more prominent as previously observed during the 16-cell stage (Maître et al., 2016). Since actomyosin contractility generates a significant portion of the surface tension of animal cells, this would translate in a gradual increase in surface tension. To investigate this, we set to measure the surface tension of cells as a readout of contractility during cleavage stages using micropipette aspiration. Contrary to our expectations, we find that surface tension gradually decreases from the zygote to 8-cell stage (Fig 4A-B) and noticeably mirrors the behavior of the period of PeCoWaCo during cleavage stages (Fig 1E). Therefore, PeCoWaCo unlikely result simply from increased contractility. Instead, the tension of blastomeres at the zygote and 2-cell stages may be too high for PeCoWaCo to become visible in most embryos. To reduce the tension of the cortex, we used low concentrations (100 nM) of the actin polymerization inhibitor Latrunculin A (Driscoll et al., 2015). Softening the cortex of 2-cell stage embryos increased the proportions of embryos displaying PeCoWaCo (Fig 4C-D, Movie 6). This suggests that PeCoWaCo become more visible thanks to the gradual softening of the cortex of blastomeres during cleavage stages. Moreover, low concentrations of Latrunculin A decreased the oscillation period of PeCoWaCo down to ~100 s, as compared to ~150 s for the DMSO control embryos (Fig 4E). This suggests that modifications of the polymerization rate of the actin cytoskeleton could be responsible for the increase in PeCoWaCo frequency observed during cleavage stages.

**Figure 4:**
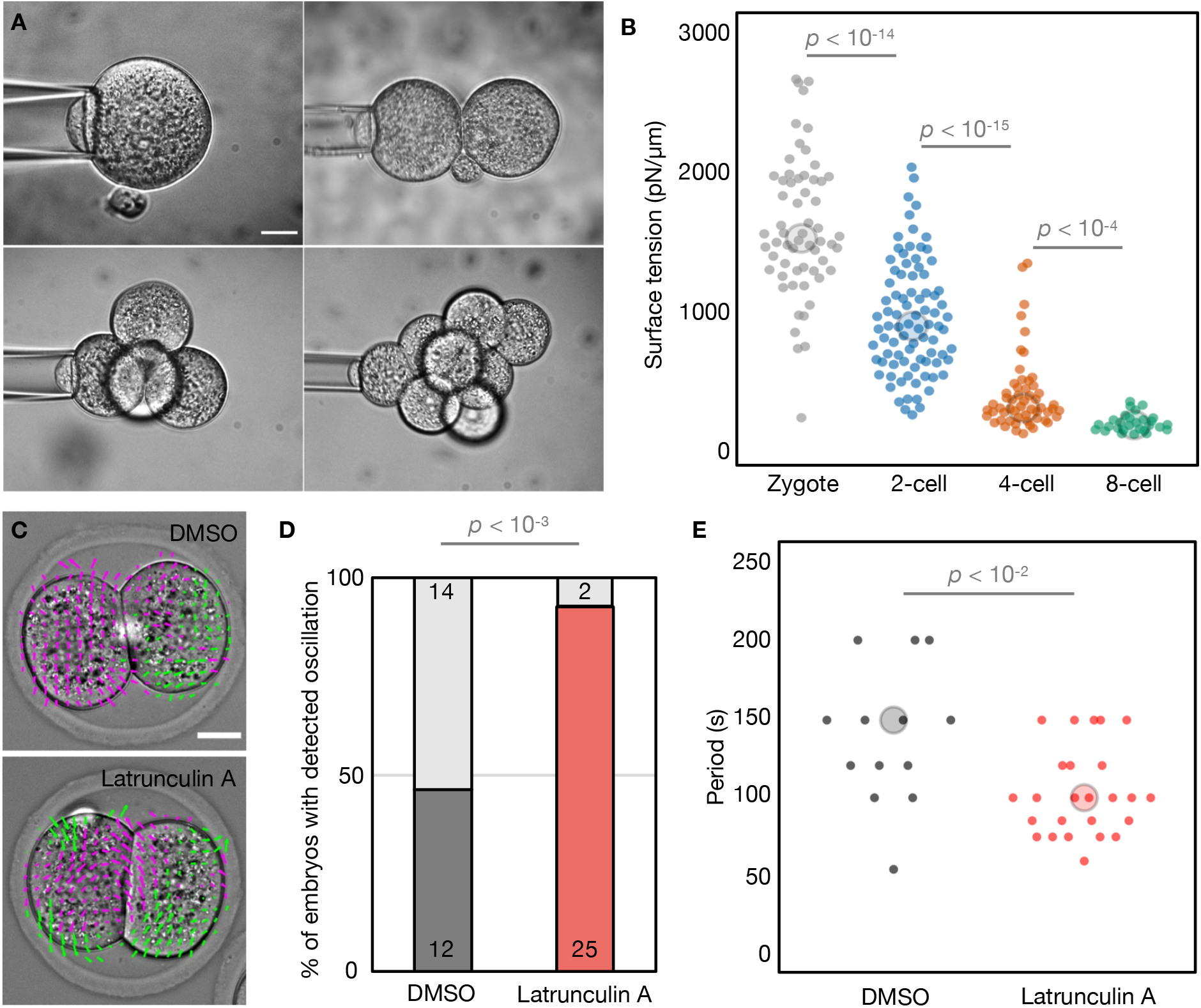
Cortical changes during preimplantation development. **A)** Representative images of the tension mapping method. **B)** Surface tension of blastomeres throughout cleavage stages. Zygote (gray, n = 60), 2-cell (blue, n = 86), 4-cell (orange, n= 55), early 8-cell (green, n=28) stages are shown. **C)** Representative images of Control and 100nM Latrunculin A treated embryos overlaid with a subset of velocity vectors from Particle Image Velocimetry (PIV) analysis (Movie 6). **D-E)** Proportion (D) of embryos showing detectable oscillations and their detected period (E) of DMSO treated (n = 26) and 100 nM Latrunculin A treated (n = 27) 2-cell stage embryos. Chi^2^ p value is indicated. Light grey shows non-oscillating embryos. Larger circles show median values.

Together, these experiments reveal the unsuspected maturation of the cortex of blastomeres during the cleavage stages of mouse embryonic development.

## Discussion

During cleavage stages, blastomeres halve their size with successive divisions. Besides the increased number of blastomeres, there is no change in the architecture of the mouse embryo until the 8-cell stage with compaction. We find that this impression of stillness is only true on a timescale of hours since, on the timescale of seconds, blastomeres display signs of actomyosin contractility. During the first two days after fertilization, contractility seems to mature by displaying more visible and frequent pulses. We further find that pulsed contractions do not rely on the successive reductions in cell size but rather on the gradual decrease in surface tension of the blastomeres. Therefore, during cleavage stages, cortical softening awakens zygotic contractility before preimplantation morphogenesis.

Previous studies on the cytoskeleton of the early mouse embryo revealed that both the microtubule and intermediate filaments networks mature during cleavage stages. Keratin intermediate filaments appear at the onset of blastocyst morphogenesis (Schwarz et al., 2015) and become preferentially inherited by prospective TE cells (Lim et al., 2020). The microtubule network is initially organized without centrioles around microtubule bridges connecting sister cells (Zenker et al., 2017). The spindle of early cleavages also organizes without centrioles similarly to during meiosis (Courtois et al., 2012; Schuh and Ellenberg, 2007). As centrioles form de novo, cells progressively transition from meiosis-like to mitosis-like divisions (Courtois et al., 2012). We find that the actomyosin cortex also matures during cleavage stages by decreasing its oscillation period (Fig 1E) and its surface tension (Fig 4B). Interestingly, this decrease in cortical tension seems to be in continuation with the maturation of the oocyte. Indeed, the surface tension of mouse oocytes decreases during their successive maturation stages (Larson et al., 2010). The softening of the oocyte cortex is associated with architectural rearrangements that are important for the cortical movement of the meiotic spindle (Chaigne et al., 2013; Chaigne et al., 2015). Therefore, similarly to the microtubule network, the zygotic actomyosin cortex awakens progressively from an egg-like state.

The maturation of zygotic contractility is likely influenced by the activation of the zygotic genome occurring partly at the late zygote stage and mainly during the 2-cell stage (Schulz and Harrison, Melissa M, 2018; Svoboda, 2018). Recent studies in frog and mouse propose that reducing cell size could accelerate zygotic genome activation (ZGA) (Chen et al., 2019; Zhu et al., 2020). We find that manipulating cell size is neither sufficient to trigger PeCoWaCo prematurely in most embryos, nor required to initiate or maintain them in a timely fashion with the expected oscillation period of the corresponding cleavage stage (Fig 2 and 3). Instead, we find that the surface of blastomeres in the cleavage stages is initially too tense to allow for PeCoWaCo to be clearly displayed (Fig 4). This is also suggested by previous studies in starfish oocytes, in which removing an elastic jelly surrounding the egg softens them and renders contractile waves more pronounced (Bischof et al., 2017). Interestingly, the changes in surface curvature caused by cortical waves of contraction may influence the signaling and cytoskeletal machinery controlling the wave (Bischof et al., 2017; Wu et al., 2018). Such mechano-chemical feedback has been proposed to regulate the period of contractions via the advection of regulators of actomyosin contractility (Munjal et al., 2015). As a result, the curvature of cells and tissues is suspected to regulate contractile waves (Bailles et al., 2019; Bischof et al., 2017). Using cell fragmentation and fusion, we have manipulated the curvature of the surface over which PeCoWaCo travel (Fig 3). From radii ranging between 10 and 30 *μ*m, we find no change in the period or travelling velocity of PeCoWaCo (Fig 3). This indicates that, in the mouse embryo, the actomyosin apparatus is robust to the changes in curvature taking place during preimplantation development. Therefore, the cleavage divisions per se are unlikely regulators of preimplantation contractility. The robustness of PeCoWaCo to changes of radii ranging from 10 to 30 *μ*m is puzzling since neither the oscillation period nor the wave velocity seem affected (Fig 3H-I). One explanation would be that the number of waves present simultaneously changes with the size of the cells. Using our 2D approach, we could not systematically analyze this parameter. Nevertheless, we did note that some portions of the fused blastomeres did not display PeCoWaCo. These may be corresponding to apical domains, which do not show prominent PeCoWaCo (Maître et al., 2016). Therefore, the relationship between the total area of the cell and the “available” or “excitable” area for PeCoWaCo may not be straightforward (Bement, 2015). In the context of the embryo, in addition to the apical domain, cell-cell contacts also downregulate actomyosin contractility and do not show prominent contractions (Maître et al., 2015). As cell-cell contacts grow during compaction and apical domains expand (Korotkevich et al., 2017; Zenker et al., 2018), the available excitable cortical area for PeCoWaCo eventually vanishes.

Together, our study uncovers the maturation of the actomyosin cortex, which softens and speeds up the rhythm of contractions during the cleavage stages of the mouse embryo. Importantly, zebrafish embryos also soften during their cleavage stages, enabling doming, the first morphogenetic movement in zebrafish (Morita et al., 2017). It will be interesting to see whether cell and tissue softening during cleavage stages is conserved in other animals. Finally, what controls the maturation of the cortex during cleavage stages will require further investigation to understand how embryos prepare for morphogenesis.

## Acknowledgements

We thank the imaging platform of the Genetics and Developmental Biology unit at the Institut Curie (PICT-IBiSA@BDD) for their outstanding support; the animal facility of the Institut Curie for their invaluable help. We thank Victoire Cachoux for help with image analysis and Ido Lavi, Sophie Louvet-Vallée, and members of the Maître lab for discussions. We thank Julie Plastino and Markus Schliffka for critical reading of the manuscript. Research in the lab of J.-L.M. is supported by the Institut Curie, the Centre National de la Recherche Scientifique (CNRS), the Institut National de la Santé Et de la Recherche Médicale (INSERM), and is funded by grants from the ATIP-Avenir program, the Fondation Schlumberger pour l’Éducation et la Recherche via the Fondation pour la Recherche Médicale, the European Research Council Starting Grant ERC-2017-StG 757557, the European Molecular Biology Organization Young Investigator program (EMBO YIP), the INSERM transversal program Human Development Cell Atlas (HuDeCA), Paris Sciences Lettres (PSL) “nouvelle equipe” and QLife (17-CONV-0005) grants and Labex DEEP (ANR-11-LABX-0044) which are part of the IDEX PSL (ANR-10-IDEX-0001-02). Ö. Ö. is funded from the European Union’s Horizon 2020 research and innovation program under the Marie Skłodowska-Curie grant agreement No 666003, the Fondation pour la Recherche Médicale (FDT202001010796), and benefitted from the EMBO YIP COVID bridging fund.

## Author contributions

Ö. Ö., L. P. and A. F. T. performed experiments and prepared data for analyses. V. K. wrote the curvature analysis code following Ö. Ö. and J.-L.M. initial plans. Ö. Ö. and J.-L.M. acquired funding, analyzed the data, designed the project and wrote the manuscript.

## Methods

### Embryo work

#### Recovery and culture

All animal work is performed in the animal facility at the Institut Curie, with permission by the institutional veterinarian overseeing the operation (APAFIS #11054-2017082914226001). The animal facilities are operated according to international animal welfare rules.

Embryos are isolated from superovulated female mice mated with male mice. Superovulation of female mice is induced by intraperitoneal injection of 5 international units (IU) pregnant mare’s serum gonadotropin (PMSG, Ceva, Syncro-part), followed by intraperitoneal injection of 5 IU human chorionic gonadotropin (hCG, MSD Animal Health, Chorulon) 44-48 hours later. Embryos are recovered at E0.5 by dissecting in 37°C FHM (LifeGlobal, ZEHP-050 or Millipore, MR-122-D) from the oviduct the ampula, from which embryos are cleared with a brief (5-10 s) exposure to 37°C hyalorunidase (Sigma, H4272).

Embryos are recovered at E1.5 or E2.5 by flushing oviducts from plugged females with 37°C FHM using a modified syringe (Acufirm, 1400 LL 23).

Embryos are handled using an aspirator tube (Sigma, A5177-5EA) equipped with a glass pipette pulled from glass micropipettes (Blaubrand intraMark or Warner Instruments). Embryos are placed in KSOM (LifeGlobal, ZEKS-050 or Millipore, MR-107-D) or FHM supplemented with 0.1 % BSA (Sigma, A3311) in 10 μL droplets covered in mineral oil (Sigma, M8410 or Acros Organics) unless stated otherwise. Embryos are cultured in an incubator with a humidified atmosphere supplemented with 5% CO2 at 37°C.

To remove the Zona Pellucida (ZP), embryos are incubated for 45-60 s in pronase (Sigma, P8811).

For imaging, embryos are placed in 5 or 10 cm glass-bottom dishes (MatTek).

Only embryos surviving the experiments were analyzed. Survival is assessed by continuation of cell division as normal when embryos are placed in optimal culture conditions.

#### Mouse lines

Mice are used from 5 weeks old on. (C57BL/6xC3H) F1 hybrid strain is used for wild-type (WT). To visualize plasma membranes, mTmG (Gt(ROSA)26Sor^tm4(ACTB-tdTomato,-EGFP)Luo^) is used (Muzumdar et al., 2007).

#### Isolation of Blastomeres

ZP-free 2-cell or 4-cell stages embryos are aspirated multiple times (typically between 3– 5 times) through a smoothened glass pipette (narrower than the embryo but broader than individual cells) until dissociation of cells.

For 16-cell-stage embryos, they are placed into EDTA containing Ca^2+^ free KSOM (Biggers et al., 2000) for 8–10 min before dissociation. Cells are then washed with KSOM for 1 h before experiment.

#### Chemical reagents and treatments

Vx-680 (Tocris, 5907) 50 mM dimethyl sulfoxide (DMSO) stock was diluted to 2.5 *μ*m in KSOM. To prevent mitosis, 2-cell-stage embryos are cultured in 2.5 *μ*m Vx-680 for 3 h shortly prior to the 2^nd^ cleavage and then washed in KSOM.

Cytochalasin D (Sigma, C2618-200UL) 10 mM DMSO stock is diluted to 10 μM in KSOM. To fragment cells, isolated 2- or 16-cell stage blastomeres were treated with Cytochalasin D for 20 min before being gently aspirated into a smoothened glass pipette of diameter about 30 or 5-10 *μ*m respectively (Korotkevich et al., 2017). 2-3 repeated aspirations are typically sufficient to clip cells into to 2 large fragments, one containing the nucleus and one without. Cells that did not fragment after 2 aspirations are used as control.

GenomONE-CF FZ SeV-E cell fusion kit (CosmoBio, ISK-CF-001-EX) is used to fuse blastomeres (Schliffka et al., 2021). HVJ envelope is resuspended following manufacturer’s instructions and diluted in FHM for use. To fuse blastomeres of embryos at the 16-cell stage, embryos are incubated in 1:50 HVJ envelope for 15 min at 37°C followed by washes in KSOM.

Latrunculin A (Tocris, ref 3973) 10mM DMSO stock is diluted to 100 nM in KSOM. To soften cells, 2-cell stage embryos are imaged in medium containing Latrunculin A covered with mineral oil for 2 h.

#### Micropipette aspiration

As described previously (Guevorkian, 2017; Maître et al., 2015), a microforged micropipette coupled to a microfluidic pump (Fluigent, MFCS EZ) is used to measure the surface tension of embryos. In brief, micropipettes of radii 8-16 μm are used to apply step-wise increasing pressures on the cell surface until reaching a deformation, which has the radius of the micropipette (*R_p_*). At steady-state, the surface tension *γ* of the cell is calculated from the Young-Laplace’s law applied between the cell and the micropipette: *γ* = *P_c_* / 2 (1/*R_p_* −1/*R*), where *P_c_* is the critical pressure used to deform the cell of radius *R*.

8-cell stage embryos are measured before compaction (all contact angles < 105°), during which surface tension would increase (Maître et al., 2015).

### Microscopy

For live imaging, embryos are placed in 5 cm glass-bottom dishes (MatTek) under a CellDiscoverer 7 (Zeiss) equipped with a 20x/0.95 objective and an ORCA-Flash 4.0 camera (C11440, Hamamatsu) or a 506 axiovert (Zeiss) camera.

Using the experiment designer tool of ZEN (Zeiss), we set up nested time-lapses in which all embryos are imaged every 5 h for 10 min with an image taken every 5 s at 2 focal planes positioned 10 μm apart. Embryos are kept in a humidified atmosphere supplied with 5% CO2 at 37°C.

mTmG embryos are imaged at the 2- and 16-cell stage using an inverted Zeiss Observer Z1 microscope with a CSU-X1 spinning disc unit (Yokogawa). Excitation is achieved using a 561 nm laser through a 63x/1.2 C Apo Korr water immersion objective. Emission is collected through 595/50 band pass filters onto an ORCA-Flash 4.0 camera (C11440, Hamamatsu). The microscope is equipped with an incubation chamber to keep the sample at 37°C and supply the atmosphere with 5% CO2.

Surface tension measurements are performed on a Leica DMI6000 B inverted microscope equipped with a 40x/0.8 DRY HC PL APO Ph2 (11506383))objective and Retina R3 camera and 0,7x lens in front of the camera. The microscope is equipped with an incubation chamber to keep the sample at 37°C and supply the atmosphere with 5% CO2.

### Data analysis

#### Image analysis

##### Manual shape measurements

FIJI (Schindelin et al., 2012) is used to measure cell, embryo, pipette sizes, and wave velocity. The circle tool is used to fit a circle onto cells, embryos and pipettes. The line tool is used to fit lines onto curvature kymographs.

##### Particle image velocimetry (PIV) analysis

To detect PeCoWaCo in phase contrast images of embryos, we use Particle Image velocimetry (PIV) analysis followed by a Fourier analysis.

As previously (Maître et al., 2015; Schliffka et al., 2021), PIVlab 2.02 running on Matlab (Thielicke and Eize J. Stamhuis, 2020; Thielicke and Stamhuis, 2010) is used to process 10 min long time lapses with images taken every 5 s using 2 successive passes through interrogation windows of 20/10 μm resulting in ∼180 vectors per embryo.

The x- and y-velocities of individual vectors from PIV analysis are used for Fourier analysis. A Fourier transform of the vector velocities over time is performed using Matlab’s fast Fourier transform function. The resulting Fourier transforms are squared to obtain individual power spectra. Squared Fourier transforms in the × and y directions of all vectors are averaged for individual embryos resulting in mean power spectra of individual embryos.

Spectra of individual embryos are checked for the presence of a distinct amplitude peak to extract the oscillation period. The peak value between 50 s and 200 s was taken as the amplitude, as this oscillation period range is detectable by our imaging method. An embryo is considered as oscillating when the amplitude peaks 1.85 times above background (taken as the mean value of the power spectrum signal of a given embryo). This threshold value was chosen to minimize false-positive and false-negative according to visual verification of time-lapse movies. For example, visual inspection of embryos shown in Fig 1D suggest that three zygote, eight 2-cell, zero 4-cell, three 8-cell are false positive, and zero zygote, two 2-cell, one 4-cell, five 8-cell are false negative. Therefore, the number of oscillating zygote is likely overestimated while the number of oscillating 8-cell stage is underestimated.

2-cell, 4-cell and 8-cell stages are considered early during the first half of the corresponding stage and late during the second half.

##### Local curvature analysis

To measure PeCoWaCo period, amplitude and velocity, we analyze the associated changes in surface curvature and perform Fourier analysis.

To obtain the local curvature of isolated blastomeres, we developed an approach similar to that of (Driscoll et al., 2012; Maître et al., 2015; Maître et al., 2016). First, a Gaussian blur is applied to mTmG images using FIJI (Schindelin et al., 2012). Then, using Ilastik (Sommer et al., 2011), pixels are associated with cell membrane or background. Segmentation of cells are then used in a custom made Fiji plugin (called *WizardofOz*, found under the *Mtrack* repository) for computing the local curvature information using the start, center and end point of a 10 μm strip on the cell surface to fit a circle. The strip is then moved by 1 pixel along the segmented cell and a new circle is fitted. This process is repeated till all the points of the cell are covered. The radius of curvature of the 10 μm strip boundaries are averaged. Kymograph of local curvature values around the perimeter over time is produced by plotting the perimeter of the strip over time.

Curvature kymographs obtained from local curvature tracking are then exported into a custom made Python script for 2D Fast Fourier Transform analysis.

Spectra of individual cells are checked for the presence of a distinct amplitude peak to extract the oscillation period. The peak value between 50 s and 200 s was taken as the amplitude, as this oscillation period range is detectable by our imaging method.

To measure the wave velocity a line is manually fitted on the curvature kymograph using FIJI.

#### Statistics

Data are plotted using Excel (Microsoft) and R-based SuperPlotsOfData tool (Goedhart, 2021). Mean, standard deviation, median, one-tailed Student’s *t*-test, Fisher exact test, and Chi^2^ *p* values are calculated using Excel (Microsoft) or R (R Foundation for Statistical Computing). Statistical significance is considered when *p* < 10^−2^.

The sample size was not predetermined and simply results from the repetition of experiments. No sample that survived the experiment, as assessed by the continuation of cell divisions, was excluded. No randomization method was used. The investigators were not blinded during experiments.

### Code availability

The code used to analyze the oscillation frequencies from PIV and local curvature analyses can be found at github.com/MechaBlasto/PeCoWaCo.git.

The Fiji plugin for local curvature analysis *WizardofOz* can be found under the *MTrack* repository.

## Movie legends

**Movie 1: Particle image velocimetry (PIV) analysis during cleavage stages.** Time-lapse imaging of zygote, 2-, 4- and 8-cell stage embryos showing PeCoWaCo. Pictures are taken every 5 s and PIV analysis is performed between two consecutive images. PIV vectors are overlaid on top of the images with vectors pointing upward in magenta and downward in green. Scale bar, 20 *μ*m.

**Movie 2: PIV analysis of 4-cell stage embryos treated with DMSO or Vx-680.** Time-lapse imaging of 4-cell stage embryos showing PeCoWaCo after treatment with DMSO or 2.5 *μ*m Vx680 at the time of the 2^nd^ cleavage division. Pictures are taken every 5 s and PIV analysis is performed between two consecutive images. PIV vectors are overlaid on top of the images with vectors pointing upward in magenta and downward in green. Scale bar, 20 *μ*m.

**Movie 3: PIV analysis of fragmented 2-cell stage blastomeres.** Time-lapse imaging of mechanically manipulated and fragmented 2-cell stage blastomeres with and without nucleus. Pictures are taken every 5 s and PIV analysis is performed between two consecutive images. PIV vectors are overlaid on top of the images with vectors pointing upward in magenta and downward in green. Scale bar, 20 *μ*m.

**Movie 4: Surface deformation tracking of fused cells.** Montage of mTmG (top) and local curvature measurements (bottom) of fused 8x, 4x, 2x 1/16^th^ blastomeres showing PeCoWaCo. Scale bar, 20 *μ*m.

**Movie 5: Surface deformation tracking of fragmented 16-cell-stage blastomeres.** Montage of mTmG (top) and local curvature measurements (bottom) of mechanically manipulated and fragmented 16-cell stage blastomeres with and without nucleus 1/16^th^ blastomeres showing PeCoWaCo. Scale bar, 10 *μ*m.

**Movie 6: PIV analysis of 2-cell stage embryos treated with DMSO or Latrunculin A.** Time-lapse imaging of 2-cell stage embryos treated with DMSO or 100 nM Latrunculin A (LatA) showing PeCoWaCo. Pictures are taken every 5 s and PIV analysis is performed between two consecutive images. PIV vectors are overlaid on top of the images with vectors pointing upward in magenta and downward in green. Scale bar, 20 *μ*m.

## Supplementary figures

**Supplementary Figure related to Figure 1:**
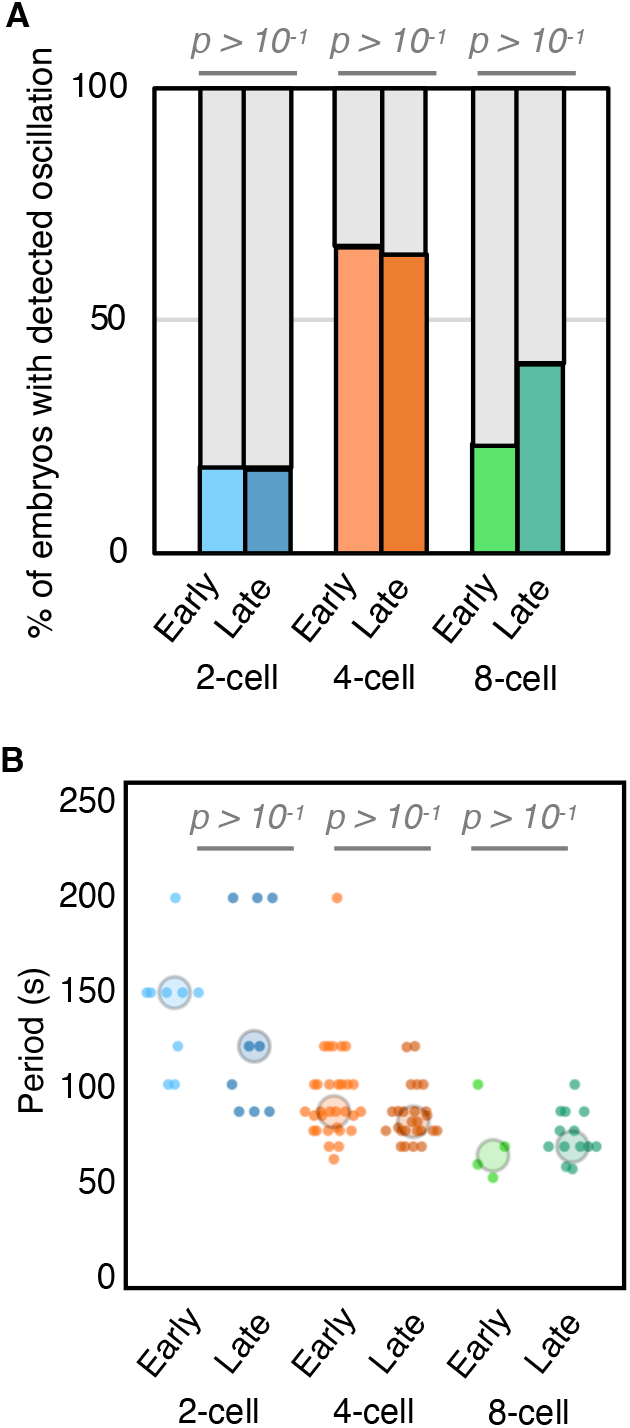
Analysis of PeCoWaCo within cleavage stages. **A)** Proportion of early and late 2-cell (blue, n = 43 and 50), 4-cell (orange, n = 38 and 28), 8-cell stage (green, n = 17 and 32) embryos showing detectable oscillations after Fourier transform of PIV analysis. Chi^2^ p values comparing different stages are indicated. Light grey shows non-oscillating embryos. **B)** Oscillation period of early and late 2-cell (blue, n = 22), 4-cell (orange, n = 33), 8-cell (green, n = 22) stages embryos. Larger circles show median values. Student’s t test p values are indicated.

**Supplementary Figure related to Figure 2:**
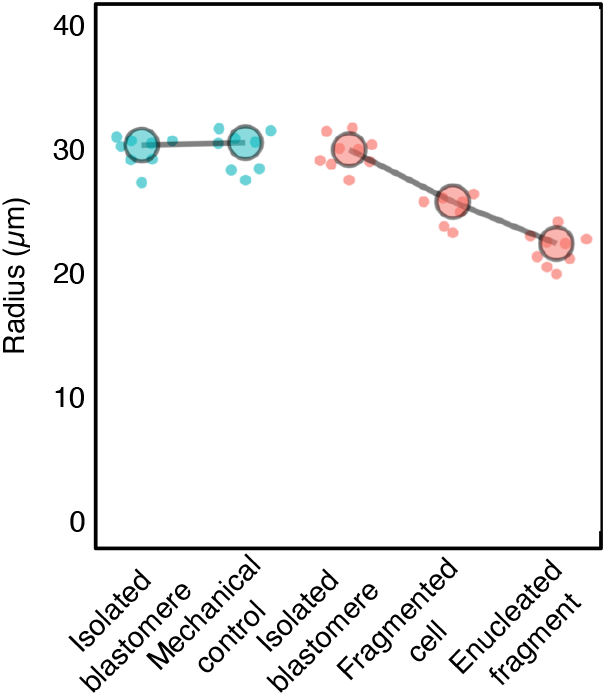
size of cells before and after fragmentation. Radius of 16-cell stage blastomeres before and after mechanical manipulation (n = 8), fragmentation into fragmented cells (n= 8) or enucleated fragments (n=9). Larger circles show median values.

